# Post-transcriptional regulation of insulin mRNA storage by G3BP1/2^+^ condensates in beta cells

**DOI:** 10.1101/2024.05.25.595863

**Authors:** Esteban Quezada, Jovana Vasiljevic, Akshaye Pal, Carla Münster, Daniela Friedland, Eyke Schöniger, Anke Sönmez, Annika Seiler, Pascal Roch, Carolin Wegbrod, Katharina Ganß, Klaus-Peter Knoch, Nicole Kipke, Simon Alberti, Rita Nano, Lorenzo Piemonti, Daniela Aust, Jürgen Weitz, Marius Distler, Michele Solimena

## Abstract

Hyperglycemia upregulates insulin translation in pancreatic beta cells. Several RNA-binding proteins involved in this process have been identified, including G3BP1, a stress granule marker downregulated in islets of subjects with type 2 diabetes. We show that in mouse insulinoma MIN6-K8 cells exposed to fasting glucose levels G3BP1 and its paralog G3BP2 colocalize to cytosolic condensates with eIF3b and *Ins1/2* mRNA. Upon glucose stimulation, the condensates dissolve and G3BP1/2, eIF3b, and insulin mRNAs redistribute throughout the cytosol. Intriguingly, G3BP1^+^ condensates in MIN6-K8 cells differ from sodium arsenate-induced stress granules in regards to eIF2α and AMPKα phosphorylation. Knockout of *G3BP1* or *G3BP2* prevented condensate assembly, but only *G3BP1* deletion decreased the levels of *Ins1/2* mRNA and proinsulin and impaired polysome formation. Like glucose, other insulin secretagogues such as Exendin-4 and palmitate, but not high KCl, prompted the dissolution of G3BP1^+^ condensates. G3BP1^+^/*Ins* mRNA^+^ condensates were also present in mouse and human beta cells from normoglycemic donors. Hence, G3BP1^+^ condensates represent a glucose-regulated compartment for the physiological storage and protection of insulin mRNA in resting beta cells.

## Introduction

Pancreatic islet beta cells play a major role in maintaining glucose homeostasis by synthesizing and secreting the peptide hormone insulin in response to elevation of blood glucose levels (Karpińska and Czauderna 2022). Insulin, in turn, lowers glycemia through insulin receptor signaling, hence promoting glucose uptake into muscle cells and adipocytes while inhibiting glycogenolysis and gluconeogenesis in hepatocytes. Accordingly, impaired beta cell function together with elevated insulin resistance cause hyperglycemia and type 2 diabetes (T2D) (Zheng et al. 2018). Beta cell dysfunction in T2D has been attributed to several mechanisms, often combined with each other, including reduced oxidative phosphorylation (Supale et al. 2012; Dludla et al. 2023), prolonged ER stress (Back and Kaufman 2012; Mustapha et al. 2021; Shrestha 2021), chronic islet inflammation (Donath et al. 2003; Donath et al. 2013; Kulkarni et al. 2022; Rohm et al. 2022) and amyloidosis (Kahn et al. 1999; Jurgens et al. 2011) resulting in beta cell dedifferentiation (Talchai et al. 2012; Son and Accili 2023) and death (Butler et al. 2003). Yet, it is important to appreciate that a clear explanation for how beta cell dysfunction evolves during the progression from normoglycemia to T2D is still missing. Also, it is conceivable that the contribution of different mechanisms to altered insulin secretion varies among different individuals, as suggested by the identification of several disease clusters (Ahlqvist et al. 2018). Recent evidence, in particular, suggests that a lower threshold for glucose-stimulated insulin release can set in motion the vicious circle between insulin resistance and hyperinsulinemia, leading eventually to beta cell decompensation and T2D (Cohrs et al. 2020; Johnson 2021).

To gain further insight into the physiology of beta cells and its pathophysiology, we undertook the systematic transcriptomic analysis of pancreatic islets retrieved by laser capture microdissection from surgical specimens of metabolically profiled living donors who underwent pancreatectomy for a variety of pancreatic disorders (Solimena et al. 2018; Barovic et al. 2019). These investigations identified the RNA-binding protein G3BP1 to be among the most significantly downregulated genes in islets of living donors with T2D compared to normoglycemic individuals (Wigger et al. 2021). G3BP1 and its paralog G3BP2, also known as Ras-GTPase Activating Protein SH3 domain-Binding Proteins 1 and 2, have been shown to be essential components of the stress granules (Guillén-Boixet et al. 2020; Yang et al. 2020; Kang et al. 2021). These non-membranous cytosolic condensates, which result from the interaction of several RNA binding proteins (RBPs) with polyadenylated mRNAs, form in cells with arrested translation under various stress conditions, such as oxidative stress, nutrient deprivation, heat, and UV radiation (Mahboubi and Stochaj 2017). Upon stressor removal, stress granules disassemble, allowing mRNAs to re-engage with ribosomes for protein synthesis (Wheeler et al. 2016). Stress granule assembly may also serve as a cellular defense mechanism in the early stages of viral infection by suspending translation (Jayabalan et al. 2021). Accordingly, some viruses have evolved strategies to disrupt stress granule functionality during later infection stages (Lloyd 2012; Yang et al. 2018). Besides G3BP1/2, other constituents of stress granules include RNA-binding proteins TIA-1/R and PABP1, the small 40S ribosomal subunit, and translation initiation factors eIF2, eIF3, eIF4A, eIF4G and eIF4E, although the composition of these condensates can vary depending on the cell type and the specific stress encountered (Moutaoufik et al. 2014; Protter and Parker 2016; Fay and Anderson 2018).

Acute elevation of insulin production in glucose-stimulated beta cells does not depend on insulin gene transcription, but on the increased translation of pre-existing copies on insulin mRNA (Itoh and Okamoto 1980; Welsh et al. 1985; Knoch et al. 2004). However, little is known about where beta cells store insulin mRNA in fasting conditions. In the present study we began to fill this gap of knowledge by investigating the possible relationship between stress granules with insulin mRNA storage sites in beta cells, as a necessary premise to understand the potential implications of downregulated gene expression of *G3BP1* in islets of subjects with T2D.

## Results

### *Ins1/2 mRNA* is enriched in G3BP1^+^/G3BP2^+^/eIF3b^+^ condensates in resting MIN6-K8 cells

Our studies began by analyzing the expression of G3BP1 and its paralogue G3BP2 in glucose-responsive mouse insulinoma MIN6-K8 cells. In these cells the mRNA levels of *G3BP1*, *G3BP2*, as well as of *Ins1* and *Ins2* did not change upon elevation of extracellular glucose from 2.8 mM (resting condition) to 16.7 mM (stimulating condition) for 30 min. (Fig. 1A). Likewise, the protein levels of G3BP1 and G3BP2 remained unchanged (Fig. 1B). Confocal microscopy showed that in resting cells G3BP1 and G3BP2 were enriched in cytosolic condensates, potentially corresponding to G3BP1^+^ and G3BP2^+^ stress granules described in other cell types (Mahboubi and Stochaj 2017). Such compartments typically form in response to conditions eliciting pronounced energy depletion, such as starvation or exposure to oxidative stress-inducing agents, to retain mRNAs while suppressing their translation, thereby reducing energy expenditure. Consistent with this possibility, G3BP1^+^/G3BP2^+^ condensates were also enriched in eIF3b, which is part of the small ribosomal unit and another marker of stress granules (Yang et al. 2020). Notably, in MIN6-K8 cells stimulated with 16.7 mM glucose, G3BP1^+^/G3BP2^+^/eIF3b^+^ were redistributed throughout the cytosol and their condensates were no longer detectable (Fig. 1C). Since stress granules also contain mRNA strands attached to RBPs and to the 40S small ribosomal subunit (Wolozin and Ivanov 2019), we examined the distribution of *Ins1/2* mRNA and the housekeeping *GAPDH* mRNA by single molecule RNA FISH (smRNA FISH). We observed that *Ins1/2* mRNA, but not *GAPDH* mRNA, was confined to G3BP1/2^+^ condensates in resting conditions (Fig. 1D). Upon stimulation with 16.7 mM glucose, all signals, including that for *Ins1/2* mRNA, diffused throughout the cells (Fig. 1E). We also addressed the distribution of mRNAs encoding for other cargoes of the insulin secretory granules, such as *Ptprn/ICA512*, *Pcsk1* and *Pcsk2,* as their translation is also rapidly upregulated in response to elevation of glucose levels to enable the biogenesis of these organelles (Knoch et al. 2004). Unexpectedly, however, none of them was detectable in G3BP1^+^ condensates of resting MIN6-K8 cells (Appendix Fig. S1).

**Figure 1.**
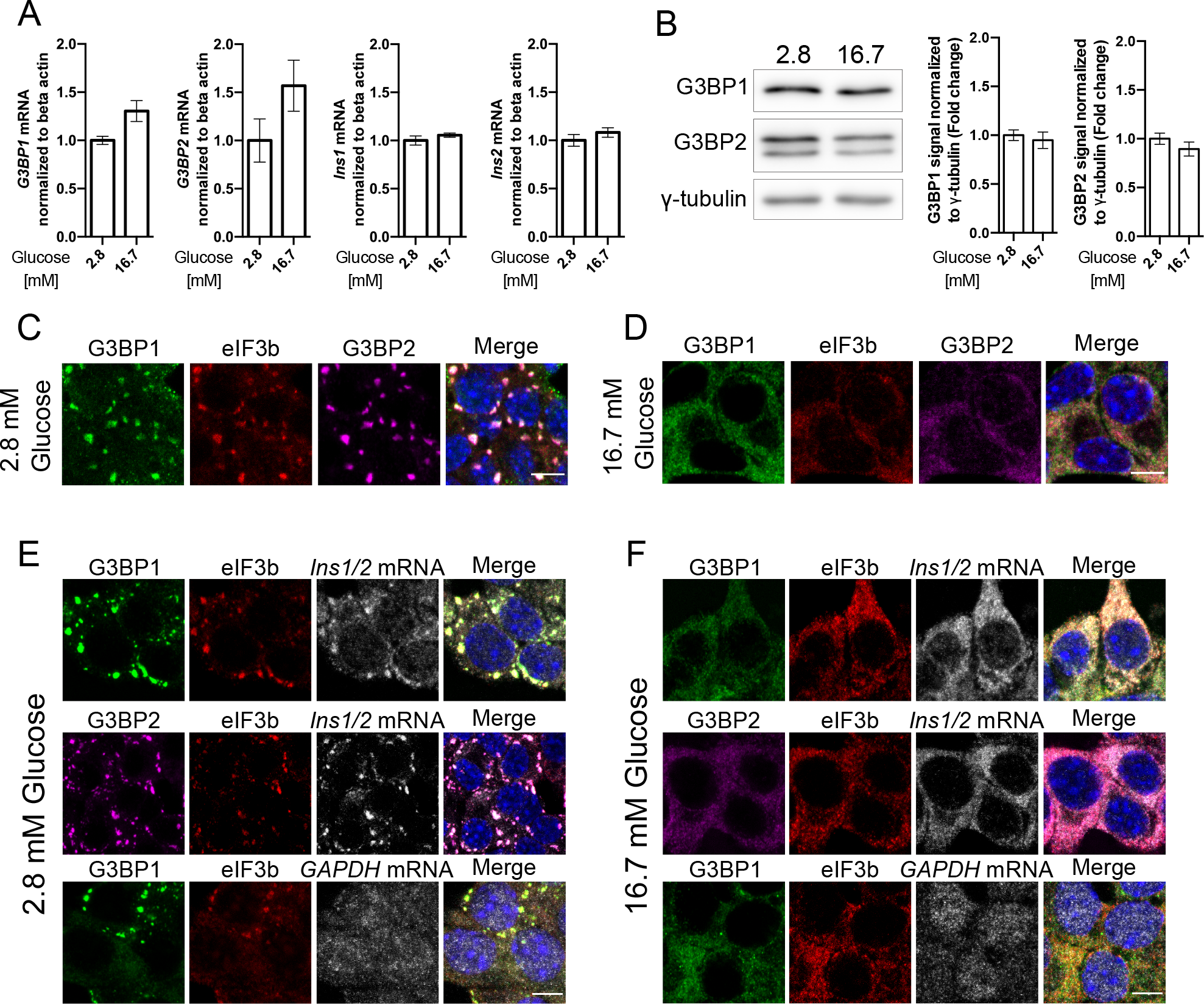
Co-localization of G3BP1, G3BP2, and eIF3b with *Ins1/2* in glucose-regulated cytosolic condensates in MIN6-K8 cells. A mRNA levels of *G3BP1*, *G3BP2*, *Ins1*, and *Ins2* in resting (2.8 mM) and stimulating (16.7 mM) glucose concentrations as measured by qRT-PCR. B Protein levels of G3BP1 and G3BP2 under resting and stimulating glucose concentrations as assessed by western blot. C Immunostaining for G3BP1 (green), G3BP2 (magenta), eIF3b (red) in resting or stimulating glucose concentrations. Nuclei are stained with DAPI (blue). D Immunostaining for G3BP1 (green), G3BP2 (magenta), eIF3b (red), and smRNA FISH for *Ins1/2* or *GAPDH* (gray) in resting glucose concentrations. Nuclei are stained with DAPI (blue). Data information: The intensity of the bands in the Western blots (B) was measured in arbitrary units using ImageStudioLite software, normalized to the γ-tubulin loading control, and fold change was calculated relative to the resting glucose condition. The values represent the mean ± s.e.m. (standard error of the mean) from three independent experiments and were analyzed via a paired *t*-test with Mann-Whitney correction. Scale bar in all imaging panels = 5 μm.

### G3BP1^+^ condensates in MIN6-K8 cells occur in conditions distinct from those associated with the biogenesis of arsenate-induced stress granules

In view of their glucose-regulated dynamics we reasoned that the G3BP1^+^ condensates observed in insulin-producing cells could reflect a physiological rather than a response to severe stress. To test this hypothesis, we examined the occurrence of G3BP1^+^ condensates in resting or glucose-stimulated MIN6-K8 cells in the presence or absence of a strong oxidative agent such as 1mM sodium arsenate, which typically elicits the formation of canonical stress granules in other cell types. As expected, glucose stimulation reduced the average number of G3BP1^+^/eIF3b^+^*/Ins1/2* mRNA^+^ condensates/cell compared to resting cells (Fig. 2A- C). In the presence of 1 mM sodium arsenate, however, the average number and size of G3BP1^+^/eIF3b^+^*/Ins1/2* mRNA^+^ condensates/cell did not change, regardless of the glucose levels (Fig. 2A-C).

**Figure 2.**
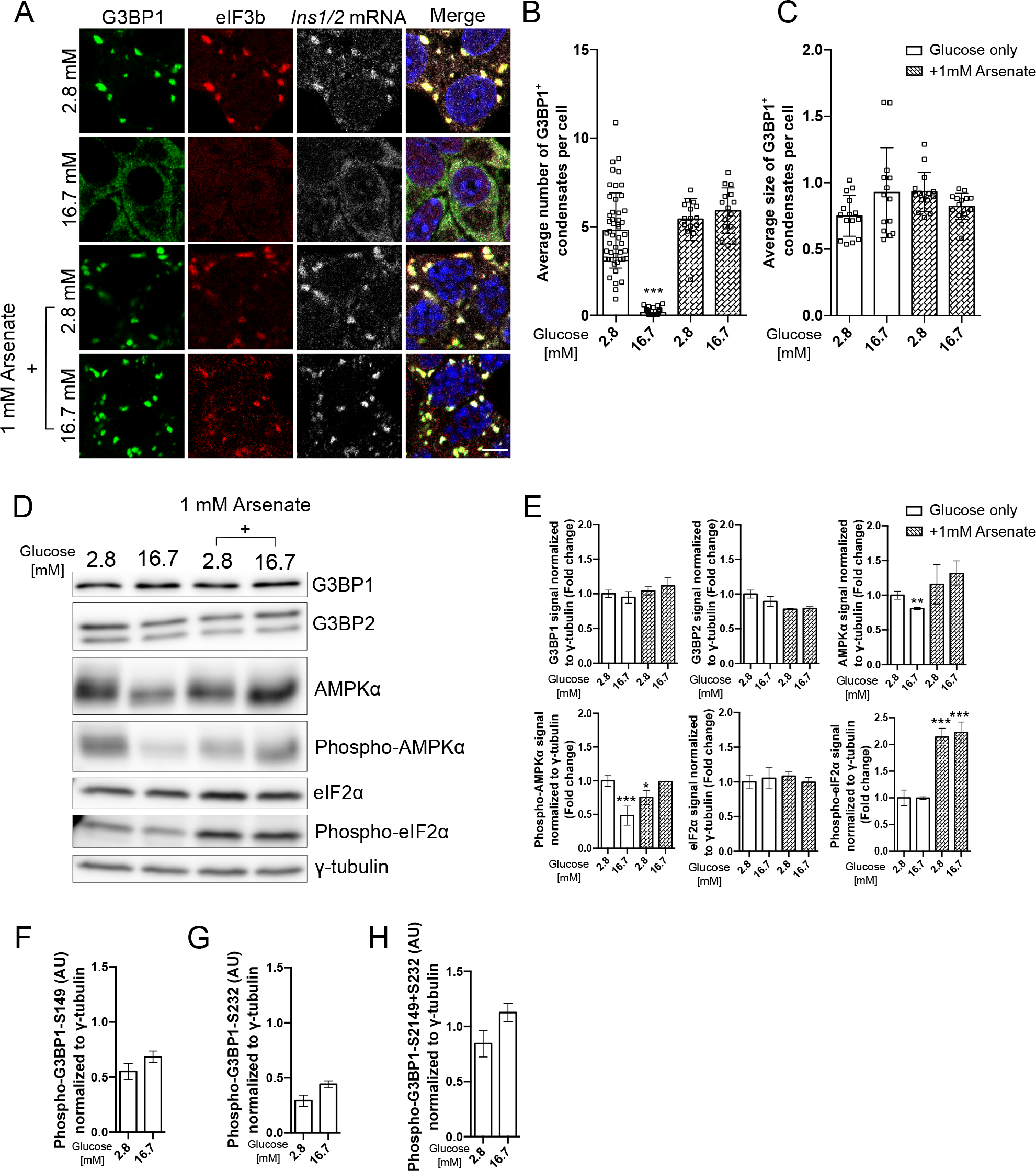
Differential behavior and signaling pathways of G3BP1^+^ condensates in response to glucose stimulation and oxidative stress in MIN6-K8 cells. A Immunostaining for G3BP1 (green), eIF3b (red) and smRNA FISH for *Ins1/2* (gray) under resting and stimulating glucose conditions with and without 1 mM sodium arsenate. Nuclei are stained with DAPI (blue). B Average number of G3BP1^+^ condensates in all tested conditions as in A. C Average size of G3BP1^+^ condensates in all tested conditions as in A. D, E Western blot (D) and quantifications (E) for G3BP1, G3BP2, AMPKα, phospho- AMPKα, eIF2α, phospho-eIF2α and gamma-tubulin across varying conditions: resting and stimulating glucose concentrations, with and without 1 mM sodium arsenate. F, G, H. Quantification of phospho-G3BP1-S149 and phospho-G3BP1-S232 alone or combined based on the Western blots shown in Fig. S3 for the corresponding phospho-G3BP1 species. Data information: The intensity of the bands in the Western blots was quantified in arbitrary units utilizing ImageStudioLite software, normalized against the γ-tubulin loading control, with fold changes for each condition calculated relative solely to the resting glucose condition. Presented values denote the mean ± s.e.m. (standard error of the mean) derived from three independent experiments, analyzed via one- way ANOVA. Values with p* < 0.05 and p*** < 0.001 were considered statistically significant relative to 2.8 mM glucose only. Scale bar = 5 μm.

In different cell types the induction of stress granules in response to various stressors involves the phosphorylation of eIF2α (Dimasi et al. 2017). On the other hand, in resting and glucose-stimulated MIN6-K8 cells treated with sodium arsenate, the levels of phospho-eIF2α were similarly elevated compared to MIN6-K8 cells exposed to low or high glucose levels alone (Fig. 2D-E). Stress granule formation has also been associated with phosphorylation of the nutrient-sensor AMPKα (Mahboubi & Stochaj, 2017). We observed a significant reduction of phospho- AMPKα levels in glucose-stimulated MIN6-K8 cells compared to resting cells (Fig. 2D-E). Unexpectedly, however, treatment of resting cells with 1 mM sodium arsenate correlated with lower rather than higher levels of phospho-AMPKα relative to cells exposed to 2.8 mM glucose only (Fig. 2D-E), while in the case of glucose-stimulated cells sodium arsenate treatment correlated with increased phospho-AMPKα levels. Studies in other cell types suggested that the assembly of stress granules is regulated through the phosphorylation of S149 and S232 within the acidic intrinsically disordered region of G3BP1 (Reineke et al. 2017; Guillén-Boixet et al. 2020; Tourrière et al. 2023). In the case of MIN6-K8 cells, glucose stimulation did not enhance the levels of G3BP1 S149 and S232 phosphorylation, either individually or cumulatively (Fig. 2F-H and Fig. S3A-C). Collectively, these results suggest that in MIN6-K8 cells G3BP1^+^/eIF3b^+^*/Ins1/2* mRNA^+^ condensation occurs in conditions that deviate from those found upon induction of stress granules by sodium arsenate, with increased phosphorylation of eIF2α, AMPKα and G3BP1, albeit in the case of the latter the relevance of its phosphorylation for stress granule biogenesis has been debated (Panas et al. 2019; Tourrière and Tazi 2019).

### G3BP1, but not G3BP2, is required for *Ins1/2* mRNA stability

G3BP1 and G3BP2 are critical constituents of stress granules. In some instances, the occurrence of stress granules requires the expression of both G3BP1 and G3BP2 (Kedersha et al. 2016; Hofmann et al. 2021), while in other cases absence of only one of these factors prevents it (Yang et al. 2020). To investigate whether G3BP1 and G3BP2 are essential components of *Ins1/2* mRNA^+^ condensates, we characterized *G3BP1^-/-^* and *G3BP2^-/-^*MIN6-K8 cells generated by CRISPR/Cas9 technology (Fig. 3A-B). Lack of either G3BP1 and G3BP2 alone was sufficient to abolish the presence of eIF3b^+^/*Ins1/2* mRNA^+^ condensates in resting cells, with the signal for both molecules being more evenly distributed throughout the cytosol (Fig. 3C-D). In the case of resting *G3BP2^-/-^*cells, G3BP1 was also more diffused compared to control cells, albeit still enriched in particles of undefined identity (Fig. 3C). Conversely, in resting *G3BP1^-/-^* cells, G3BP2 was homogeneously distributed in the cytoplasm (Fig. 3D). In *G3BP1^-/-^* cells these phenotypic alterations were associated with reduced levels of both insulin transcripts, and especially *Ins1* mRNA, regardless whether the cells were resting (Fig. 3E) or had been harvested from the culture media with 25 mM glucose (Fig. S2A). In *G3BP2^-/-^*cells *Ins1* mRNA levels were unchanged relative to wild-type cells, while those of *Ins2* mRNA were elevated independently of the glucose concentration (Fig. 3C and Fig. S2A). Accordingly, G3BP1 seems to exert a protective/stabilizing role for *Ins1/2* transcripts, whereas G3BP2 may be implicated in *Ins2* mRNA degradation. At the protein level, *G3BP1^-/-^* cells displayed a lower proinsulin content in both resting and glucose-stimulated conditions compared to wild-type cells (Fig. 3F-G), although the amount of insulin was unchanged (Fig. 3H). Conversely, in *G3BP2^-/-^* cells neither proinsulin nor insulin content were altered compared to wild-type cells (Fig. 3F-G).

**Figure 3.**
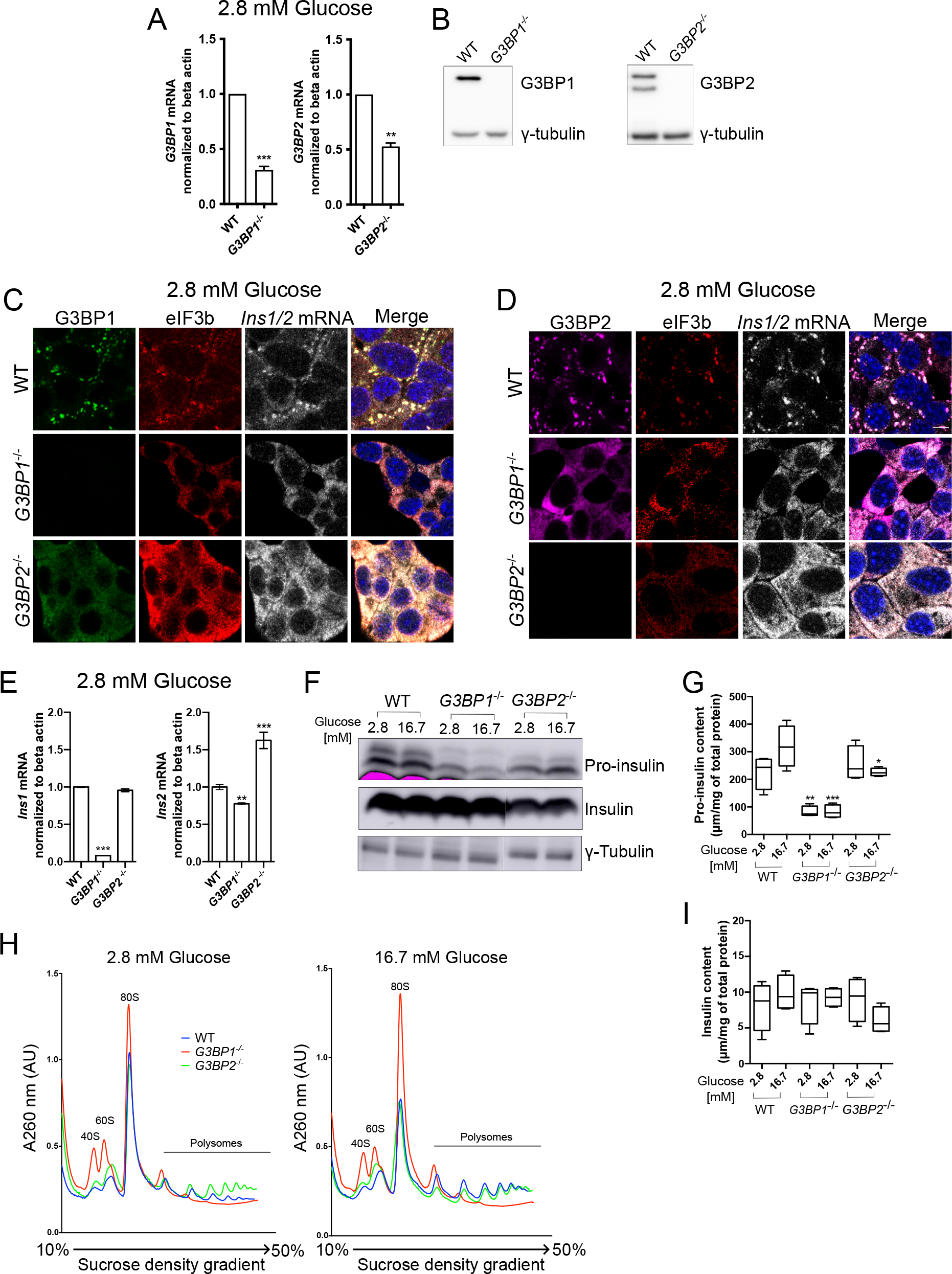
G3BP1, unlike G3BP2, safeguards *Ins1/2* mRNA, ensures apt prepro- insulin translation, and modulates the polysome turnover in response to glucose variations. A mRNA levels of *G3BP1* and *G3BP2* in wild type (WT) *G3BP1^-/-^* and *G3BP2^-/-^* MIN6-K8 cells as measured by qRT-PCR. B Protein levels of G3BP1 and G3BP2 and gamma-tubulin in WT and *G3BP1^-/-^* and *G3BP2^-/-^* MIN6-K8 cells as measured by western blot. C, D Immunostaining for G3BP1 (green), eIF3b (red), G3BP2 (magenta) and smRNA FISH for *Ins1/2* in WT, *G3BP1***^-/-^** and *G3BP2***^-/-^**MIN6-K8 cells at resting glucose concentrations. Scale bars = 5 µM E Quantification of *Ins1* and *Ins2* in WT, *G3BP1***^-/-^**and *G3BP2***^-/-^** cells in resting glucose conditions as assessed by qRT-PCR. F Western blotting for proinsulin, insulin and gamma-tubulin in WT, *G3BP1***^-/-^** and *G3BP2***^-/-^** in glucose resting or stimulated MIN6-K8 cells. G and I Quantification of proinsulin and insulin levels as measured by ELISA and HTRF, respectively, in WT, *G3BP1***^-/-^** and *G3BP2***^-/-^** cells under resting and stimulating glucose concentrations. H Polysomal profiling of resting or glucose-stimulated WT, *G3BP1^-/-^*and *G3BP2^-/-^* MIN6-K8 cells using a 10%-50% sucrose gradient to separate and enrich in fractions of the small ribosomal units (40S, first peak), the big ribosomal units (60S, second peak), the monosomes (80S, third peak) and polysome subpopulations starting with 2 ribosomes per polysome (fourth peak) until 7 ribosomes per polysome (last peak). Data information: Presented values denote the mean ± s.e.m. (standard error of the mean) derived from three independent experiments, analyzed via one-way ANOVA. Values with p* < 0.05, p** < 0.01, and p*** < 0.001 were considered statistically significant relative to the WT condition. Scale bar = 5 μm.

It has been shown that G3BP1 plays a role in polysome turnover (Meyer et al. 2020). To ascertain if this could account for lower proinsulin levels in *G3BP1^-/-^* cells, a polysomal profiling was conducted using a 10%-50% sucrose gradient. This analysis revealed that wild-type cells responded aptly to varying glucose levels, containing a pronounced monosomal peak (80S) in resting conditions (Fig. 3I), which decreased by half upon glucose stimulation, when polysome peaks were enriched (Fig. 3K). This suggests that in stimulating conditions the cells transition to active protein translation, as evidenced by the enrichment of polysomal subpopulations peaks. Instead, in *G3BP1^-/-^* cells only the first two polysomal subpopulation peaks were present (Fig. 3I), while upon glucose stimulation the height of the 80S peak was twice the size relative to wild-type cells (Fig. 3K). This profile suggests that absence of G3BP1 does not enable translation, as monosomes cannot transition into polysomes. The polysomal profile of *G3BP2^-/-^* cells, on the other hand, resembled that of wild-type cells in both resting and glucose stimulating conditions (Fig. 3K).

### Incretin and palmitate resolve G3BP1^+^ condensates

Numerous stimuli foster insulin secretion and/or synthesis (Nauck et al. 2021). Among them are the GLP-1 analogue Exendin-4 (Göke et al. 1993; Alarcon et al. 2006; Gandasi et al. 2018) and palmitate (Usui et al. 2019). Accordingly, treatments with either 50 nM Exendin-4 or 50 μM palmitate reduced G3BP1^+^/eIF3b^+^*/Ins1/2* mRNA^+^ condensates in resting MIN6-K8 cells, though not to the extent observed upon stimulation with high glucose alone (Fig. 4A-B). On the contrary, more, albeit smaller, G3BP1^+^/eIF3b^+^*/Ins1/2* mRNA^+^ condensates were observed in cells incubated with 55 mM KCl (Fig. 4A-B), which triggers the secretion of insulin without enhancing its synthesis (Hatlapatka et al. 2009). Glucokinase, or hexokinase IV, catalyzes the conversion of glucose into glucose-6-phosphate in pancreatic beta cells, thus it serves as a glucose sensor for insulin secretion due to its high flux control coefficient on glucose metabolism (Thilagavathi et al. 2022). In resting MIN6- K8 cells treated with 3 µM glucokinase activator Ro-28-1675 G3BP1^+^/eIF3b^+^*/Ins1/2* mRNA^+^ condensates were completely absent, unlike in cells exposed to DMSO alone, indicating that activation of glycolysis, even in the presence of low glucose levels, triggers the resolution of G3BP1^+^/eIF3b^+^*/Ins1/2* mRNA^+^ condensates (Fig. 4B).

**Figure 4.**
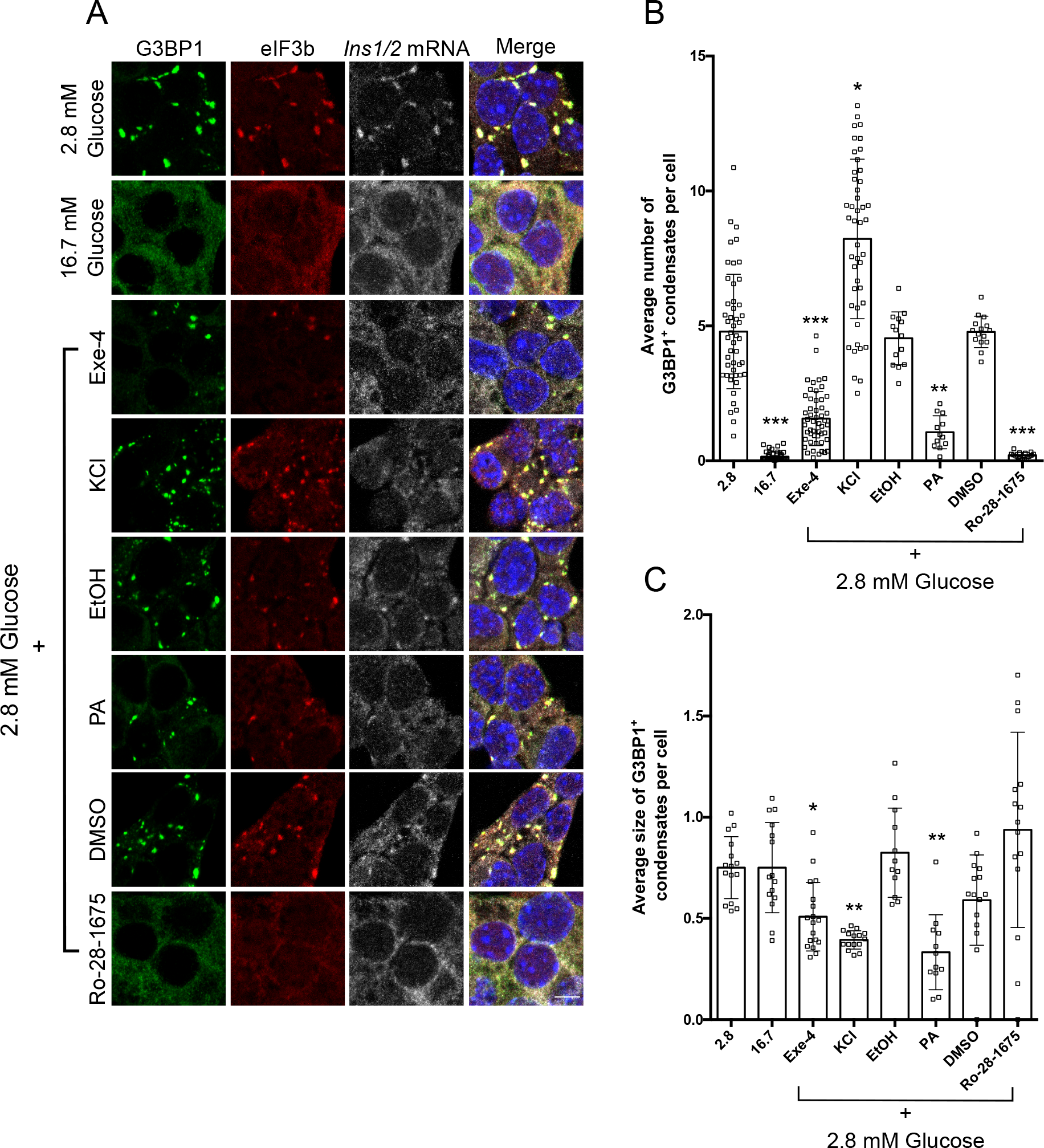
Impact of various insulin secretagogues on G3BP1^+^ condensates A Immunostaining for G3BP1 (green), eIF3b (red) and smRNA FISH for *Ins1/2* (gray) in MIN6-K8 wild-type cells treated with Exendin-4 (50 nM), palmitate (50 μM), KCl (55 mM) and glucokinase activator Ro-28-1675 (3 μM) in resting glucose conditions. B, C Quantification of number and size of G3BP1^+^ condensates in MIN6-K8 cells in resting glucose concentrations upon addition of Exendin-4 (50 nM), palmitate (50 mM), KCl (55 mM) or Ro-28-1675 (3 μM). Data information: Presented values denote the mean ± s.e.m. (standard error of the mean) derived from three independent experiments, analyzed via one-way ANOVA. Values with p* < 0.05, p** < 0.01, and p*** < 0.001 were considered statistically significant relative to the WT condition. Scale bar = 5 μm.

### G3BP1^+^/ and G3BP2^+^/*Ins1/2* mRNA^+^ condensates are present in resting mouse primary beta cells

To verify the physiological relevance of our findings in MIN6-K8 cells, we investigated the occurrence of G3BP1/2^+^ condensates in primary beta cells and their relationship with *Ins1/2* mRNA. In dispersed mouse pancreatic islet cells exposed to 2.5 or 3.3 mM glucose, both G3BP1^+^ and G3BP2^+^ condensates were present in beta cells, as identified by co-immunostaining for insulin, and enriched in *Ins1/2* mRNA (Fig. 5A-B). Conversely, these condensates were absent in dispersed islet cells incubated with 16.7 mM glucose, in which the signals for both proteins and *Ins1/2* mRNA were more diffused in the cytoplasm (Fig. 5A-B). To preclude the possibility of artifacts arising from the dispersion of pancreatic islets, similar analyses were conducted on whole mount mouse pancreatic islets. Consistent with the observations made in dispersed islet cells, resting beta cells within isolated islets also displayed G3BP1^+^/*Ins1/2*^+^ mRNA condensates (Fig. 5C-D). Therefore, the phenotype observed within the mouse insulinoma MIN6-K8 cells was validated in primary mouse beta cells.

**Figure 5.**
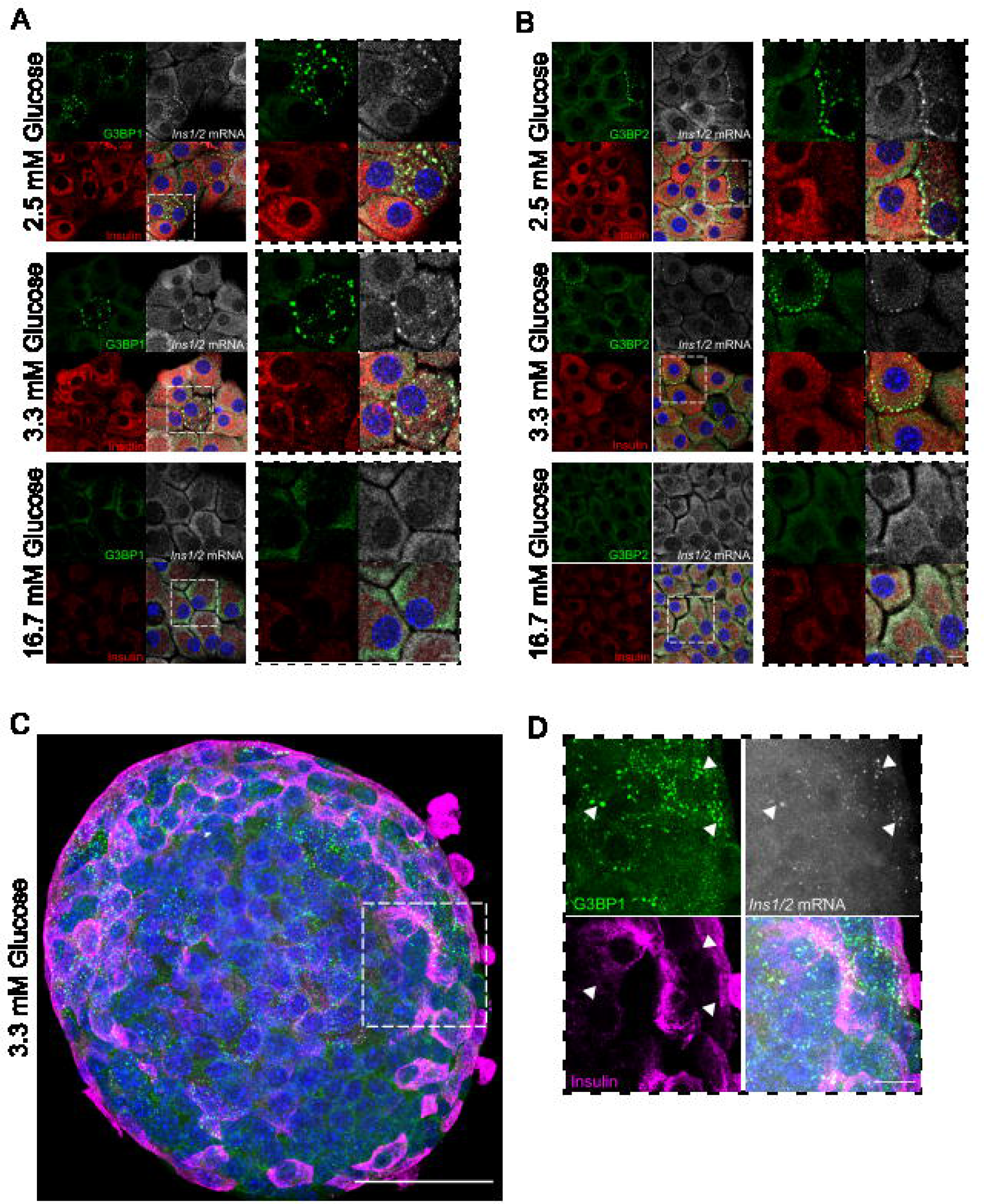
Detection of G3BP1^+^, G3BP2^+^ and *Ins1/2* in dispersed and whole mouse pancreatic islet cells under resting glucose conditions. A, B Immunostaining for G3BP1 (green, left panel), G3BP2 (green, right panel), Insulin (red) and smRNA FISH for *Ins1/2* (gray) in dispersed mouse pancreatic islet cells under glucose resting (2.5 and 3.3 mM) or stimulating (16.7 mM) conditions. Nuclei are stained with DAPI (blue). C, D Immunostaining for G3BP1 (green), Insulin (magenta) and smRNA FISH for *Ins1/2* (gray) in whole mount mouse pancreatic islets under resting conditions. Arrowheads point to G3BP1^+^/*Ins1/2* mRNA^+^ condensates. Nuclei are stained with DAPI (blue). Data information: Scale bars in A, B and D = 5 μm; in C = 50 μm.

### *INS* mRNA^+^ condensates are present in pancreatic islet beta cells in situ of non-diabetic living donors

Transcriptomic profiling of laser capture microdissected islets from metabolically phenotyped human living donors in our LIDOPACO cohort had revealed that *G3BP1* is downregulated in individuals with type 2 diabetes compared to non-diabetic donors (Wigger et al. 2021). Hence, we investigated whether G3BP1^+^/*INS* mRNA^+^ condensates could also be detected in beta cells within snap-frozen surgical specimens of living donors undergoing pancreatectomy for a variety of disorders of the exocrine pancreas. Initial attempts to identify such condensates through a random survey of samples from normoglycemic living donors (NLD) were unfruitful. This was not unexpected, as the threshold for stimulation of insulin translation and secretion of human beta cells is ∼4 mM glucose, i.e. lower than in rodents, while even at fast the average glycemia of normoglycemic subjects in our cohort (65) was 5.03 ± 0.32 mM. For this reason, we took advantage of the continuous monitoring of glycemia during surgery to select among the NLD within our cohort those with the lowest intraoperative glycemia in the minutes before pancreas resectomy. Intriguingly, in specimens of two such donors, coded NLD1 and NLD2, several pancreatic islets displayed *INS* mRNA^+^ condensates, which especially in the case of NLD1, partially colocalized with G3BP1 (Fig. 6). This pattern, however, was not observed in specimens of other non-diabetic donors with higher intraoperative glycemia (Fig. 6). Nonetheless, these findings document the occurrence of G3BP1^+^/*INS* mRNA^+^ condensates in human beta cells in situ, pointing to the physiological significance of such dynamic compartments for control of insulin translation.

**Figure 6.**
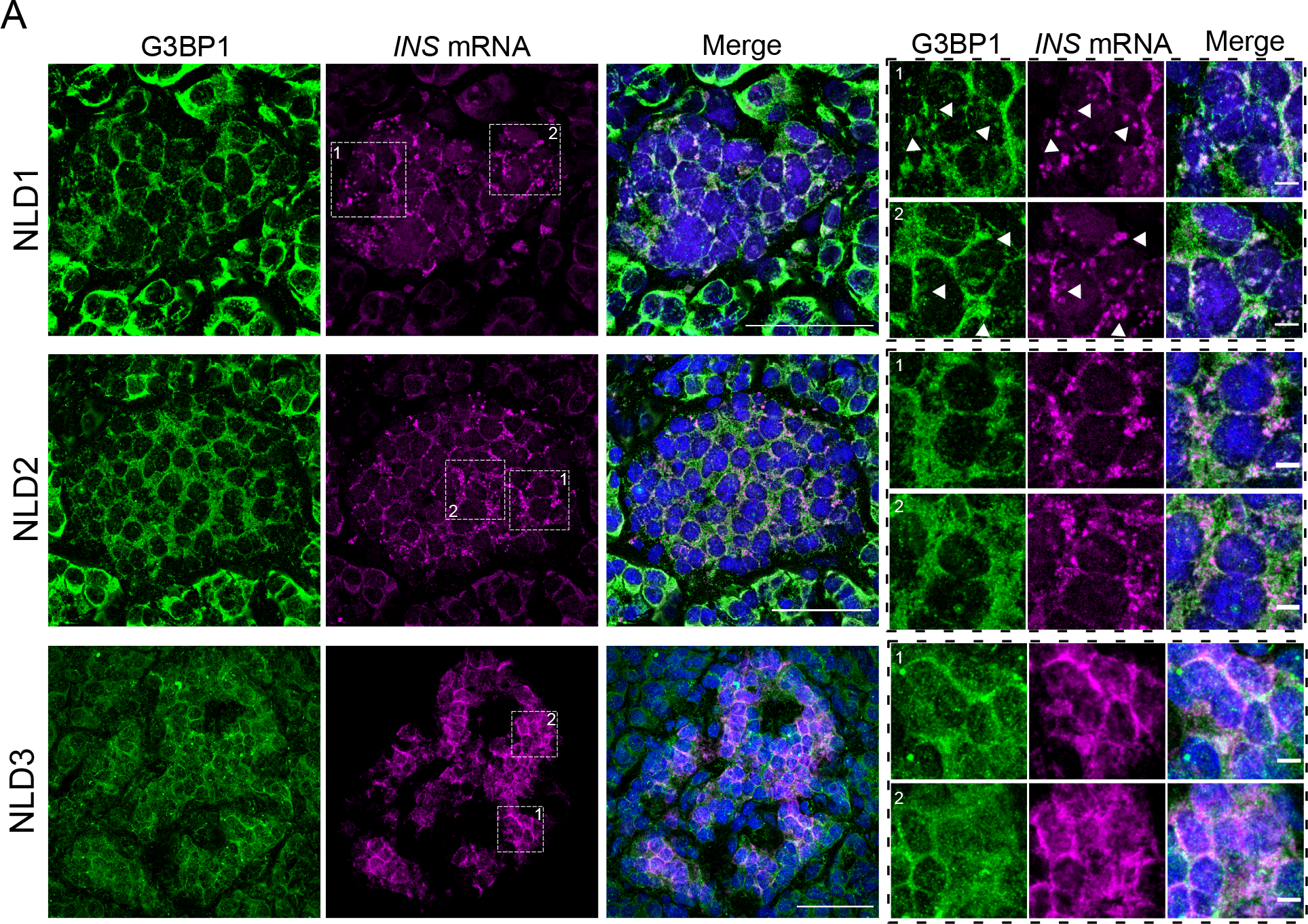
G3BP1^+^ and *INS* mRNA^+^ condensates in beta cells in situ of normoglycemic living donors. A Immunostaining for G3BP1 (green), and smRNA FISH for *INS* (magenta) in human pancreatic cryosections from normoglycemic living donors (NLD) 1, 2 and 3. Nuclei are stained with DAPI (blue). For better visualization, the right panels show magnified crops of the area selected with white line squares in the left panels. Arrowheads point to G3BP1^+^/*INS* mRNA^+^ condensates. Data information: Scale bars = 50 μm or 5 μm in the left and right panels, respectively.

## Discussion

In this study, we unveil the presence of G3BP1/2^+^ and eIF3b^+^ cytosolic condensates, which encapsulate *Ins1/2* mRNAs at resting glucose concentrations and dissolve when beta cells are exposed to higher glucose levels. Intriguingly, neither the housekeeping *GAPDH* mRNA nor mRNAs encoding for other glucose-induced insulin secretory cargoes *Ptprn*, *Pcsk1*, *Pcsk2* were found within G3BP1^+^ condensates, suggesting the potentially selective confinement of *Ins1/2* mRNA in such novel beta cell compartment. As stress granules can capture various mRNAs, including housekeeping ones like *GAPDH* mRNA (Kedersha and Anderson 2002), the condensates described here may not be equivalent to conventional stress granules induced by stress conditions. However, *Ins1/2* transcripts are by far the most abundant in beta cells, which might account for their accumulation and detectability into G3BP1^+^ condensates.

By synthesizing and secreting insulin when exposed to 16.7 mM glucose, but not when resting with 2.8 mM glucose (Iwasaki et al. 2010), MIN6-K8 cells are a well suited model to study the dynamics of G3BP1^+^ condensates. Due to their occurrence in the presence of 2.8 mM glucose, G3BP1^+^ condensates cannot be readily attributed to stress resulting from glucose deprivation. However, since MIN6-K8 cells are cultured in media with 25 mM glucose, the drastic change to resting glucose levels could represent a potential stress. Therefore, we tested the effect of sodium arsenate, a known inducer of stress granules. In its presence, G3BP1^+^ condensates persisted even in the presence of 16.7 mM glucose. Notably, G3BP1^+^ condensates resolved rapidly, i.e. within 30 minutes, following glucose stimulation, in contrast to canonical stress granules, which in HeLa cells require >2 hours to disappear upon removal of sodium arsenate (Tolay and Buchberger 2021). Hence, beta cells may use a similar machinery to stress granules for physiological purposes.

Stress granules may mainly form as a consequence of polysome disassembly, which can be elicited in several ways: phosphorylation of eIF2α, inhibition of mTOR or interference with the eIF4F complex (Hofmann et al. 2021). mTOR inhibition can be promoted by phosphorylation of AMPK-α on T172 by LKB1/AXIN (Lin and Hardie 2018). This event occurs in low glucose concentrations when the glycolytic enzyme aldolase, unbound to its substrate fructose 1,6 biphosphate (FBP), recruits AXIN to v-ATPase, Ragulator, and AMPK-α, forming a supramolecular complex on lysosome surfaces, leading to LKB1-mediated AMPK-α phosphorylation. Concurrently, AMPK- α is phosphorylated through the canonical pathway due to an increase in the AMP/ADP ratio resulting from low glucose, which in turn decreases glycolytic flux. We observed that in low glucose conditions, G3BP1^+^ condensates appear and AMPK-α phosphorylation is elevated in comparison to cells exposed to high glucose levels, consistent with findings that AMPK-α levels are rapidly reduced, plausibly due to degradation upon glucose stimulation. Furthermore, a non-canonical pathway of stress granule formation involving AMPK-α phosphorylation induced by energy depletion has been discovered in HeLa cells, where AMPK-α interacts directly with G3BP1 (Mahboubi & Stochaj, 2017). Based on this literature and our findings, there seems to be a connection in insulinoma MIN6-K8 cells between low glucose, AMPK- α phosphorylation/activation, and presence of G3BP1^+^ condensates, possibly via the interaction of phospho-AMPK-α with G3BP1.

Despite both *G3BP1^-/-^* and *G3BP2^-/-^*MIN6-K8 cells lacking the ability to form condensates, we found contrasting effects on *Ins1/2* transcript variants and proinsulin levels when compared to wild-type cells. In *G3BP1^-/-^* cells, unlike in *G3BP2^-/-^* cells, both *Ins* transcripts and proinsulin levels were decreased. Rasputin, the Drosophila homolog of G3BP1, has been shown to bind and stabilize short mRNAs implicated in transcription, splicing, and translation during embryogenesis, thereby enhancing their translation (Laver et al. 2020). Additionally, expression of a dominant-negative variant of G3BP1 lacking the NTF2L domain, and consequently unable to dimerize or form condensates, activated intra-axonal mRNA translation and axon growth in rat dorsal root ganglia (DRG) neurons, hence expediting nerve regeneration *in vivo* (Sahoo et al. 2018). This data underscores a role for G3BP1 beyond stress responses, highlighting its function in physiological conditions where it may bolster mRNA stability and translation. Correspondingly, reduction of *Ins1/2* transcript and proinsulin levels in *G3BP1^-/-^* cells only indicates that G3BP1, but not its close paralogue G3BP2, play a role in stabilizing and enhancing the translation of insulin mRNAs.

Previous studies suggested that both *G3BP1* and *G3BP2* are required for stress granule assembly (Kedersha et al. 2016; Yang et al. 2020). In our case, deletion of either *G3BP1* or *G3BP2* alone were sufficient to elicit this phenotype. *G3BP1^-/-^* MIN6- K8 cells carry a 25 bp deletion in *G3BP1* exon 2, potentially resulting in the expression of a truncated mutant encompassing 56 residues in total, 23 of which belong to the G3BP1 NTF2L domain (Figure S4A). Whether such putative peptide may still dimerize with G3BP2 and thereby inhibit stress granule formation cannot be excluded. Notably, it was shown that truncation of USP10, a G3BP1 binding protein which is also essential for stress granule formation (Takahashi et al. 2013), generated a peptide which precludes the occurrence of stress granules (Kedersha et al. 2016). In the case of *G3BP2^-/-^* MIN6-K8 cells, the 12 bp deletion in exon 2 results in the removal of residues 36-39 (Figure S4B), which according to AlphaFold Structure Prediction, lie in the first beta sheet of the protein, likely affecting its folding and stability. We remark that in the aforementioned studies stress granules were induced by pharmacological agents triggering eIF2α phosphorylation (sodium arsenate, clotrimazole, thapsigargin) or eIF4A inactivation (pateamine-A, rocaglamide A), and not by modulating glucose concentrations within a range mimicking physiological condition.

G3BP1 has also been identified as a key player in the ribosome-associated quality control, overseeing the fidelity of mRNA translation, particularly when aberrant translation occurs (Meyer et al. 2020). Beta cells have an outstanding translation performance, as they can translate >3 x 10^3^ preproinsulin molecules/second/beta cell (Schuit et al. 1991; Vasiljević et al. 2020), hence necessitating a highly efficient surveillance mechanism. Translation can be compromised by degradation via non-stop decay (NSD) (Van Hoof et al. 2002; Vasudevan et al. 2002; Meyer et al. 2020), or in no-go decay (NGD) (Harigaya and Parker 2010; Shoemaker and Green 2012; Meyer et al. 2020). Both NSD and NGD are triggered by collisions of ribosomes, prompting their disassembly (Simms et al. 2017; Juszkiewicz et al. 2018; Meyer et al. 2020). In such instances, G3BP1 forms complexes with the G3BP1-family member USP10 to deubiquitinate 40S ribosomal proteins, safeguarding them from lysosomal degradation and maintaining ribosomal subunit balance (Meyer et al. 2020). In the absence of the protective deubiquitination activity of G3BP1-family- USP10 complexes, the 40S proteins along with *Ins1/2* transcripts may be degraded.

The lack of a dynamic polysomal response in glucose-stimulated G*3BP1^-/-^* MIN6-K8 cells is consistent with this scenario, and thus accounts for their reduced proinsulin translation and content.

Glucose is key for insulin secretion and biosynthesis (Leibiger et al. 2000). Yet, other molecules, such as the incretin hormone glucagon-like peptide-1 (GLP-1), potentiates glucose-stimulated insulin production and release (Thorens 1992; Alarcon et al. 2006; Nauck et al. 2021). Accordingly, exposure of resting MIN6-K8 cells to the GLP1-analogue Exendin-4 reduced the number and size of G3BP1^+^ condensates, pointing to a priming effect of GLP-1 on beta cell responsiveness to elevation of glycemia (Göke et al. 1993). This finding is particularly interesting, considering previous literature stating that increased glycolytic flux, where aldolase is substrate-bound, inactivates AMPKα (Lin and Hardie 2018). Free fatty acids (FFAs), such as palmitate, induce insulin secretion from isolated human pancreatic islets exposed to physiological fasting glucose levels, a process linked to increased glycolytic flux and mitochondrial respiration (Cen et al. 2016). This mechanism might explain the observed reduction in G3BP1^+^ condensates with the addition of palmitate to resting glucose levels. Similar considerations could apply to the reduction of G3BP1^+^ condensates upon treatment with the allosteric glucokinase activator Ro-28- 1675 (Grimsby et al. 2003), which enhances glycolysis. Exposure of beta cells to high levels of KCl is commonly used to induce membrane depolarization and stimulate insulin secretion (Brüning et al. 2017). Unlike glucose, however, this treatment does not promote insulin production (Hatlapatka et al. 2009). Accordingly, KCl treatment did not induce the loss of G3BP1^+^ condensates, but rather an increase in their number. Sulfonylureas, the oldest class of medications for the treatment of type 2 diabetes, akin to KCl treatment, induce membrane depolarization and insulin secretion by binding to the sulfonylurea receptor subunit on ATP-sensitive potassium channels (Costello et al. 2024). Despite their insulin secretagogue activity, these drugs are now considered secondary line treatments due to their long-term detrimental effects on beta cell viability, as they stimulate the release of insulin, but not its biosynthesis (Mohan et al. 2022). Furthermore, sulfonylureas have been linked to endoplasmic reticulum stress (Qian et al. 2008), and increased ROS production, leading to oxidative stress (Tsubouchi et al. 2004). Consequently, the higher prevalence of G3BP1^+^ condensates in KCl-treated cells compared to treatment with resting glucose alone could be indicative of a stress response.

Having access to pancreatic tissue from a large cohort of normoglycemic living donors who underwent pancreatic surgery enabled us to detect cytosolic *INS* mRNA^+^granular structures that partially co-localized with G3BP1^+^ condensate-like compartments, albeit not as distinctly as seen in mouse beta cells. This observation is particularly noteworthy considering the complexity of the physiological and clinical context. Albeit at fast preoperatively, these patients indeed receive various treatments during the long surgical procedure of pancreatectomy, including possibly the infusion of glucose and insulin, hence creating conditions that significantly differ from the controlled environments of our previous experiments *in vitro*.

## Materials and Methods

### Mice

Animal husbandry was done in accordance with applicable animal welfare legislation (e.g. RL 2010/63/EU). Experiments were licensed by the respective authorities in Sachsen. Mice were housed in individually ventilated cages in a 12-hour light and 12-hour dark cycle with food and water *ad libitum*.

### Human living donors and pancreatic tissue

Pancreatic surgical tissue from the “healthy” resection margins was obtained from living donors who underwent pancreatectomy at our University Hospital due to various disorders of the exocrine pancreas. All donors provided their informed consent for the use of surgical and blood samples for research purposes and our studies were conducted with the ethical approval of the Ethical Committee of the Technische Universität Dresden under the license EK151062008. Donors underwent an oral glucose tolerance test in the days immediately preceding pancreatectomy and their glycemia was monitored during surgery. After excision, the pancreatic tissue was immediately snap-frozen, then embedded in OCT and stored at -80 **°**C. Cryosections of 10 μm were generated with a cryostat (Epredia™ CryoStar™ NX50 Kryostat, Fisher Scientific) for further analysis.

### Culture of MIN6-K8 cells

MIN6-K8 insulinoma cells (Iwasaki et al., 2010) were cultured in DMEM High-Glucose (25mM) medium (ThermoFisher, 11965092) supplemented with 15% fetal bovine serum (FBS) (A5258701, gibco), 1% penicillin-streptomycin and 70 mM 2- Mercaptoethanol at 37°C in a humidified atmosphere with 5% CO2. The culture media was sterilized using a Rapid Flow Filter Unit (ThermoFisher, 566-0020). For cell expansion, 175 cm² flask bottles (Corning, 353112) were used, and cells were subsequently seeded in various plates depending on the specific techniques being employed further in the experiments.

### Treatments of MIN6-K8 cells with different glucose buffers

MIN6-K8 insulinoma cells grown in 25 mM glucose were first equilibrated to a resting state with 2.8 mM glucose at 37 °C for 30 minutes, and then either exposed without washing to a stimulation buffer with 16.7 mM glucose or kept in fresh resting glucose buffer at 37 °C for 30 minutes. Resting glucose buffer was composed of 15 mM Hepes pH 7.4, 55 mM KCl, 2.8 mM Glucose, 60 mM NaCl, 24 mM NaHCO_3_, 1 mM MgCl_2_, 2 mM CaCl_2_, 1mg/ml bovine serum albumin (Roth, 8076.3). Stimulation glucose buffer was composed of 15 mM Hepes pH 7.4, 55 mM KCl, 25 mM Glucose, 60 mM NaCl, 24 mM NaHCO_3_, 1 mM MgCl_2_, 2 mM CaCl_2_, 1mg/ml bovine serum albumin (Roth, 8076.3). Both buffers were sterile filtered using Rapid Flow Filter Unit and kept at 4°C. Buffers were warmed in a water bath at 37°C for at least 15 minutes before performing the treatments. MIN6-K8 cell culture media was washed twice with PBS 1X before adding the resting glucose buffer. After the treatment cells were processed in different ways depending on the desired technique to be performed downstream.

### Treatments of MIN6-K8 cells with different insulin secretagogues

MIN6-K8 cells were treated with various reagents to assess stress responses and activation mechanisms, using the protocols outlined in the methods. Sodium hydrogen arsenate heptahydrate (Alfa Aesar, 33373.14) at a concentration of 1 mM was added to either resting or stimulation glucose buffers to induce oxidative stress. Palmitic acid (Sigma, P-5585) was prepared as a 10 mM stock in absolute ethanol and used at 50 μM, Exendin-4 (Sigma, E7144) was prepared as a 50 mM stock in autoclaved water and used at 50 nM, Ro-28-1675 (MedChemExpress, HY-10595) was kept as a 10 mM stock in DMSO and used at 3 μM, and KCl was used at 55 mM; all these reagents were added exclusively to resting glucose buffer. Each treatment was conducted in parallel with controls that included the respective vehicle in the resting buffer.

### Protein extraction and quantification

MIN6-K8 cells were harvested from 6-well plates at a confluence of 80-90%. Cells were scraped from each well and pelleted in Eppendorf tubes in the centrifuge at 4°C x 5.000 rpm for 2 minutes in PBS 1X. Then, PBS was removed and 30-50 μL of lysis were added depending on if the pellet was from 1 or 2 wells. For quantifying total protein concentration, the Pierce™ BCA Protein Assay Kit (ThermoFisher, 23225) was used as described by the manufacturer. Samples were kept at -80°C for long term storage.

### SDS-PAGE and Western blots

Protein samples (20-30μg) were subjected to SDS-PAGE (Sodium Dodecyl Sulfate-Polyacrylamide Gel Electrophoresis) for separation based on molecular weight. Proteins were first denatured for 5 min at 95°C. The samples were then loaded onto a polyacrylamide gel in specific % depending on the target protein and electrophoresis was carried out at a constant 180 volts for 2 hours. Following electrophoresis, proteins were transferred from the gel onto a nitrocellulose membrane in a transfer buffer in a wet transfer system for 2 hours at 400 mA. After the transfer, membranes were blocked with a solution of 5% fat free dehydrated milk in PBST 0.1%. Subsequently, the membrane was incubated with horseradish peroxidase (HRP)-conjugated secondary antibodies in a dilution of 1:6.000 in PBST- T 0.1% and 5% fat free dehydrated milk. Protein bands were visualized by treating the membrane with SuperSignal™ West Pico PLUS Chemiluminescent Substrate (ThermoFisher, 34580) as described in the kit protocol and then images were taking using the Amersham Imager 600. See Table EV1 for further details about gel %, primary antibodies dilution and catalog numbers.

For lambda phosphatase assay the reactive from Lambda Protein Phosphatase (Lambda PP) (P0753L, NEB) were used and followed the protocol from the manufacturer.

### RNA Extraction, cDNA preparation and Quantitative PCR (qPCR)

MIN6-K8 cells were harvested from 6-well plates at a confluence of 80-90%. Cells were scrapped from each well and pelleted in Eppendorf tubes in the centrifuge at 4°C x 5.000 rpm for 2 minutes in PBS 1x. RNA extraction from cell pellet was performed by using RNeasy kit (Qiagen, 74104) following manufacturers protocol. The concentration of total RNA (ng/ml) per sample was measured with Nanodrop TM1000 spectrophotometer from Thermo.

For the synthesis of complementary DNA (cDNA), a total of 1μg of total RNA was used from each sample. Reactions were prepared using the M-MLV Reverse Transcriptase protocol (Promega, M1705) as described by the manufacturer. Enzyme was added to the reaction 10 minutes after incubating all the components at 72°C, to avoid heat inactivation. Then, for the reverse transcription reaction samples were incubated for 40 minutes at 42°C and then last incubation was for 10 minutes at 70°C. After the reaction was done, 1 volume of water was added to the final reaction (40μl final volume).

For quantitative PCR reactions (qPCR), reagents from GoTaq® qPCR Mastermix (Promega, A6002) were used. Reactions were prepared in technical duplicates and based in manufacturer’s instructions using 1 μl of cDNA for each reaction. qPCR reactions were conducted in 96-well plates in the AriaMx Real-Time PCR System from Agilent. Genes evaluated and their respective primer sequences can be found in Table EV2. Fold changes between conditions were calculated based on the deltadeltaCt method as described by (Livak and Schmittgen 2001).

### Immunohistochemistry and RNA fluorescent *in situ* hybridization (RNA-FISH)

For performing immunohistochemistry only, MIN6-K8 cells at 70-80% of confluence grown on 12 mm glass coverslips (Roth, P231.1) in 24-well plates (Corning, 353047) were fixed with 4% PFA (Santa Cruz, sc-281692) for 20 minutes under the chemical hood. Fixed cells were immediately stained or stored at 4°C for up to 2 weeks in PBS-Azide 0.02%. Fixed cells were permeabilized with PBS-Triton 0.1% for 15. Then, cells were blocked with blocking solution (0.2% Fish skin gelatin [Sigma, G7765] and 0.5% BSA, sterile filtered) for 30 minutes in a humidified chamber. Next, cells were incubated in primary antibody for 30 minutes in a humidified chamber. Specific antibodies used and their respective dilutions can be found in Table EV3. Next, secondary antibody incubation in a dilution of 1:1.000 for 30 minutes in a humidified chamber. DAPI nuclear staining (Sigma, D9564) was either included in the secondary antibody solution at a dilution of 1:5.000 or after this incubation DAPI was prepared in PBS 1X at a dilution of 1:5.000-1:10.000 and cells were incubated for 5 minutes. Coverslips were mounted using mowiol (Calbiochem, 475904) on glass slides and air dried at room temperature protected from light until the next day. Next day the glass slides were either imaged in the microscope or stored at 4°C for short- and long-term storage.

For Immunohistochemistry coupled with smRNA FISH, same procedure as described for immunohistochemistry only, except the addition of DAPI which is included at the end of the smRNA FISH technique. After secondary antibody incubation, cells were quickly washed 3-4 times with PBS 1X and then fixed again with PFA 4% for 10 minutes. Then, cells were washed 3 times with PBS 1X. Next, for smRNA FISH we used Stellaris technology, whether commercial ready to use probes for *Ins1/2* transcripts (Biosearch Technologies, VSMF- 3612-5) or manually designed assays for *PTRPN*, *Pcsk1* and *Pcsk2* transcripts. First, 300μL of buffer A were added to each well of a 24-well plate and incubated for 5 minutes at room temperature. Then, probes were prepared in hybridization buffer (HB [biotechne, SMF-HB1]) by adding 1μl of probe to 1 mL of hybridization buffer (buffer had 900μl of hybridization buffer and 100μl of formamide [Roth, P040]). Cells were incubated in HB overnight at 37°C in a parafilm-sealed humidified chamber. Next day, cells were washed once with buffer A for 30 minutes at 37°C. Then, buffer A was removed, and next buffer A was added again but this time including DAPI in a dilution of 1:10.000 and again incubated for 30 minutes at 37°C. Then, buffer A was removed and now buffer B was added for 5 minutes at room temperature. Finally, excess of buffer B was removed with a paper towel and then coverslips were mounted using mowiol on glass slides. Glass slides were set to air dry at room temperature and protected from light overnight. Next day the glass slides were either imaged in the microscope or stored at 4°C for short- and long-term storage.

### CRISPR-Cas9 Mediated Gene Knockout

CRISPR-Cas9 system was utilized to generate G3BP1 and G3BP2 single knockout MIN6-K8 cell lines. Guidelines published to perform this technique were followed exactly as reported in (Ran et al. 2013). To generate the sgRNAs, the online tool from Integrated DNA Technologies (IDT).

After transfections with the plasmids were performed, cells were taken to the Fluorescence- Activated Cell Sorting (FACS) facility to sort them based on GFP signal, sorting 1 cell per well in 96-well plates.

### Insulin and Pro-insulin quantification by HTRF and ELISA

Cells were treated with resting and stimulation buffers as described before. After treatment, buffer was separated in 3 aliquots and stored at -20°C. Cells were harvested and pelleted as described before. Cell pellets were resuspended in lysis buffer and followed the same protocol for protein extraction mentioned before. Total protein extraction was also separated into 3 aliquots. For measuring insulin levels, the Insulin Ultra-Sensitive Assay kit (cisbio, 62IN2PEG) was performed as described by the manufacturer. For measuring pro-insulin levels, the Rat/Mouse Proinsulin ELISA (Mercodia, 10-1232-01) was performed as described by the manufacturer.

### Polysome profiling

Polysome profiling was performed as described in the article from (Aboulhouda et al. 2017). Briefly, cell lysates were prepared by using specific reagents and buffer to ensure the preservation of ribosomes and mRNA integrity. Then, cell lysates were loaded onto sucrose gradients going from 10% to 50% of sucrose density and after the gradients were ultracentrifuged for 2 hours and 30 minutes, to separate the polysomes based on their size, with heavier complexes migrating further. Finally, sucrose gradients were fractionated in an automated way by using the Piston Gradient Fractionator from Biocomp. A chart was obtained based on the absorbance at 260 nm of the sample revealing peaks for distinct components, such as 40S, 60S, 80S and polysomes. Collected fractions were snap frozen with liquid nitrogen and then stored at -80°C.

### Mouse dispersed pancreatic islets and whole mount mouse pancreatic islets

Pancreatic islets were obtained from wild-type mice of the inbred C57JBL/6J strain. The protocol to prepare dispersed mouse pancreatic islets was adapted from (Phelps et al. 2017). Briefly, protocol begins with sterilizing and ECM-coating cover slips. Mouse islets are washed with PBS and digested using a mixture of Accutase and Cell Dissociation Solution at 37°C. The digestion is halted with ice-cold FBS. The islets are then gently dispersed into a single-cell suspension and seeded onto the prepared cover slips. Following initial incubation at 37°C, additional media is added. The cells are left undisturbed for several days to facilitate adhesion and spreading prior to subsequent experiments.

### Immunohistochemistry and smRNA FISH in mouse dispersed or whole pancreatic islets

Immunohistochemistry coupled with smRNA FISH was performed with the same protocol mentioned before for MIN6-K8 cells. A few details are different for dispersed and whole mount pancreatic islets, the glass coverslips used were of 24 x 24 mm (Roth, KCY2.1) and placed in 6-well plates (Corning, 353046). In addition, for the whole mount mouse pancreatic islets, permeabilization was performed with PBS-Triton 0.3% for 20 minutes.

### Immunohistochemistry and RNA scope in frozen human pancreatic tissue

Sections 10 μm human pancreatic cryosections were let to air dry for 5 minutes at room temperature, then a hydrophobic barrier was made by outlining the pancreatic tissue. Tissue was fixed for 15 minutes with PFA 4%, then 2 quick washes were performed with PBS 1X. Then, for INS RNA detection the protocol for RNAscope® Multiplex Fluorescent Reagent Kit v2 described by the manufacturer for frozen tissue sections was used. Subsequently, the tissue was again fixed with PFA 4% and washed twice with PBS 1X. Then, tissue was blocked with a drop of Dako antibody diluent (S0809, Dako) for 30 minutes. Then primary antibody for G3BP1 (abcam, ab181150) was prepared in Dako antibody diluent in a 1:400 dilution. Tissue was washed 3 times with PBS 1X, then secondary antibody was prepared in Dako antibody diluent in a dilution of 1:1.000 and including DAPI in a dilution of 1:10.000. Tissue was then washed 3 times with PBS 1X. Finally, 20 μl of mowiol were added on top of the tissue and then a 22 x 22 mm glass coverslip was placed on top of the mowiol drop trying to avoid bubble formation. Glass slides with the mounted coverslips were left to air dry at room temperature overnight. Next day the glass slides were either imaged in the microscope or stored at 4°C for short- and long-term storage.

### Statistical analysis

Number of independent experiments are mentioned in each figure legend. All the statistical analyses were performed with GraphPad Prism Software. Tests such as Student’s t-test and one-way ANOVA were indicated in each figure legend. Significant differences were depicted with stars for each graph and explained in the figure legends.

## Acknowledgements

We thank Janani Natarajan for helpful discussion, Mrs. Katja Pfriem for administrative support. Work in the Solimena was supported by the German Center for Diabetes Research (DZD e.V.), which is financed by the German Ministry for Education and Research; the DFG Grant SO 818/10-1; the European Union Funded Network “INTERCEPT-T2D” 101095433; the Innovative Medicines Initiative 2 Joint Undertaking under grant agreement 115881 (RHAPSODY), which includes financial contributions from the European Union’s Framework Program Horizon 2020, the European Federation of Pharmacological Industries and Associations (EFPIA), and the Swiss State Secretariat for Education, Research and Innovation under contract 16.0097. Views and opinions expressed are however those of the authors only and don’t necessarily reflect those of the funding agencies. Neither the European Union nor any of the granting authorities can be held responsible for them.

## Author contributions

Conceptualization, E.Q., J.V. and M.S.; methodology, E.Q., and J.V.; validation, E.Q., and J.V.; formal analysis, E.Q., J.V., A.P., and M.S.; investigation, E.Q., P.R., A.S., K.G., C.M., D.F., E.S., A.S., C.W., K-P.K., and N.K.; resources, M.D., D.A., J.W., R.N., and L.P.; writing – original draft, E.Q.; writing – review & editing, E.Q. and M.S.; visualization, E.Q.; supervision, S.A., and M.S.; funding acquisition, J.V. and M.S.

---

Figure S1. Immunostainings for G3BP1 (green) and smRNA FISH for *Ptprn*, *Pcsk1* and *Pcsk2* (gray) in MIN6-K8 cells in resting glucose conditions.

Figure S2. mRNA and protein levels of insulin secretory granule cargoes in MIN6-K8 wild type, *G3BP1^-/-^* and *G3BP2^-/-^*cells.

A *Ins1*, *Ins2*, *Ptprn*, *Pcsk1* and *Pcsk2* mRNA levels in WT, *G3BP1*^-/-^ and *G3BP2*^-/-^ MIN6-K8 cells, as assessed by qRT-PCR.

B-D Western blot and relative quantifications for ICA512/PTPRN, PC1/3, PC2 species in glucose resting and stimulated WT, *G3BP1*^-/-^ and *G3BP2*^-/-^ MIN6-K8 ^cells^.

E-G Respective quantifications of western blots in B and D for Pro-ICA512, ICA512 Transmembrane Fragment (ICA512-TMF), Pro-PC1/3, PC1/3, Pro-PC2 and PC2.

Data information: Presented values denote the mean ± s.e.m. (standard error of the mean) derived from three independent experiments for qRT-PCR and five independent experiments for western blots, analyzed via one-way ANOVA. Values p** < 0.01, and p*** < 0.001 were considered statistically significant relative to the WT condition.

Figure S3. Western blots for phospho-G3BP1 S149 and phospho-G3BP1 S149.

A-C Western blot for Phospho-G3BP1 S149, Phospho-G3BP1 S232 and gamma-tubulin on extracts of MIN6-K8 cells in growth media with 25 mM glucose and treated with or without lambda phosphatase.

B Western blot for Phospho-G3BP1 S149, Phospho-G3BP1 S232 and gamma-tubulin on extracts of glucose resting or stimulated MIN6-K8 cells.

C, D Quantification of the western blots shown in panel A.

Data information: Presented values denote the mean ± s.e.m. (standard error of the mean) derived from three independent experiments, analyzed via paired *t*-test with Mann-Whitney correction. Values with p*** < 0.001 were considered statistically significant relative to the -lambda condition.

Figure S4. Characterization of G3BP1 and G3BP2 deletion in *G3BP1^-/-^* and *G3BP2^-/-^* MIN6-K8 clones.

A, B Schematic illustrations of the G3BP1 (A) and G3BP2 (B) domain and exon structures. The exons coding for the NTF2L domain, which is responsible for G3BP1 and G3BP2 dimerization are colored in green. The location of the nucleotide deletions identified in *G3BP1*^-/-^ and *G3BP2*^-/-^ MIN6-K8 clones and the alterations in the corresponding amino sequences are shown. The deletion in *G3BP1* introduces a premature stop codon. The deletion in *G3BP2* removes four amino acids and converts Y40 into N.

## Tables

Table 1. Antibodies list

Table 3. Blood glucose levels of normoglycemic living donors

Table 2. Primer sequences

## Notes

### Competing Interest Statement

The authors have declared no competing interest.

