## Supplementary figures and images for "Post-transcriptional regulation of insulin mRNA storage by G3BP1/2^+^ condensates in beta cells"

### Figure S1

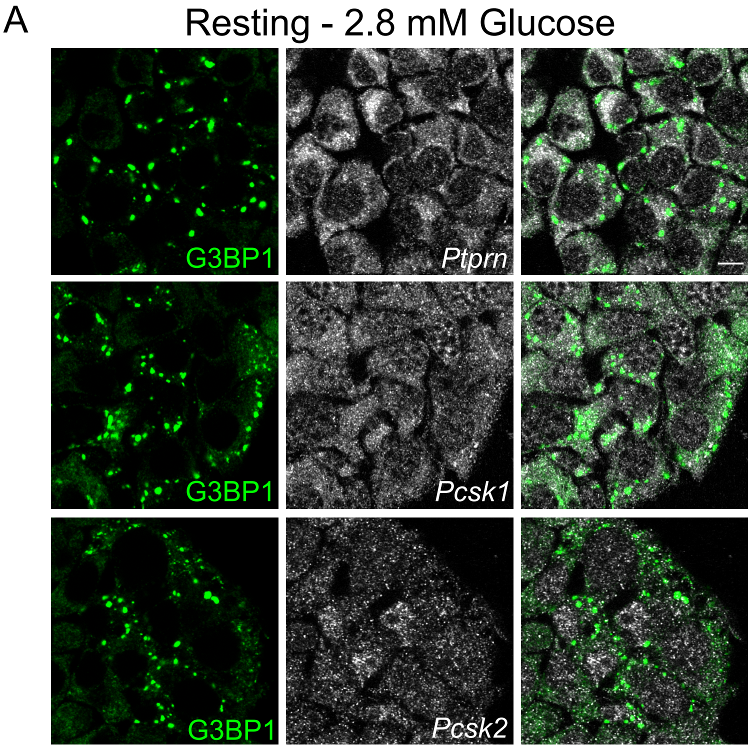

### Figure S2

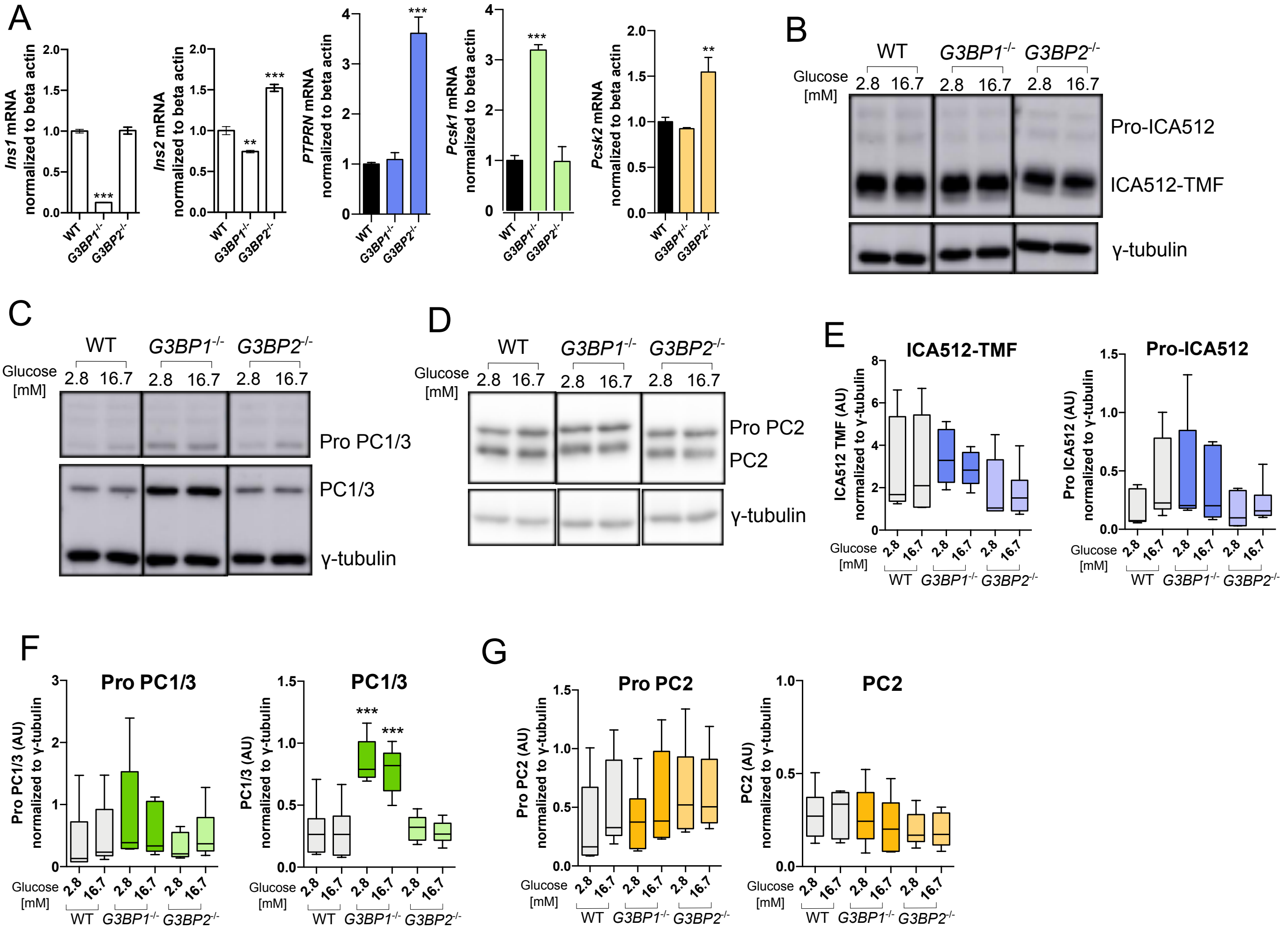

### Figure S3

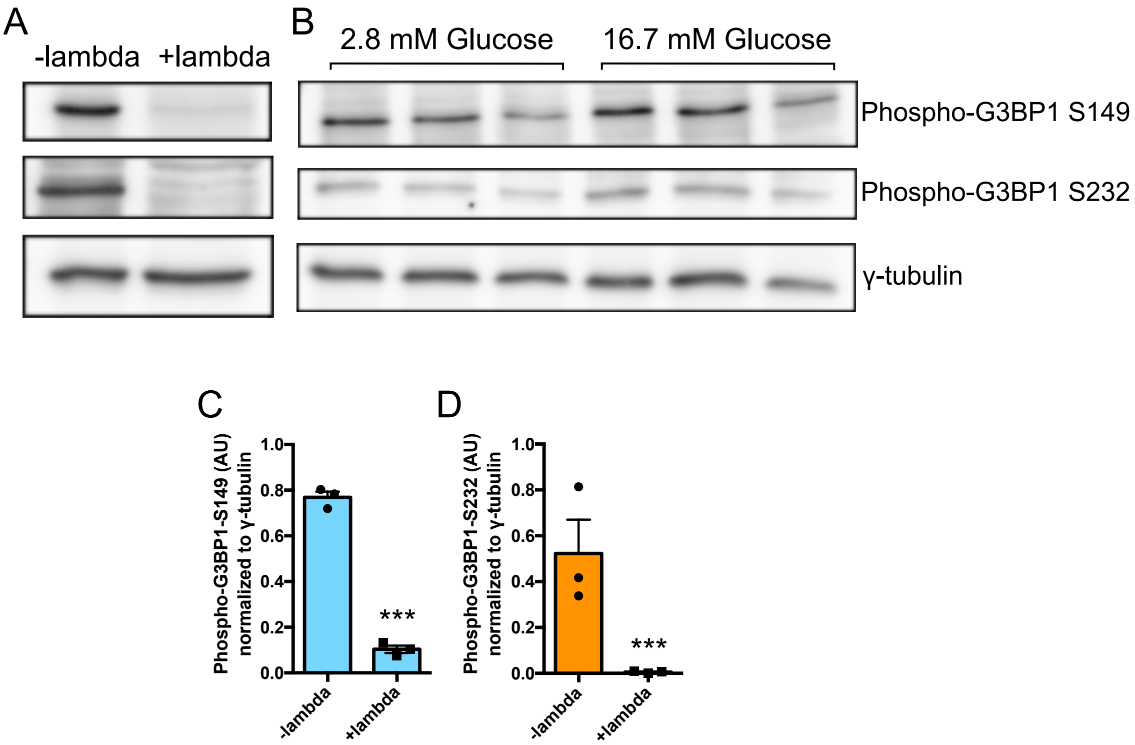

### Figure S4

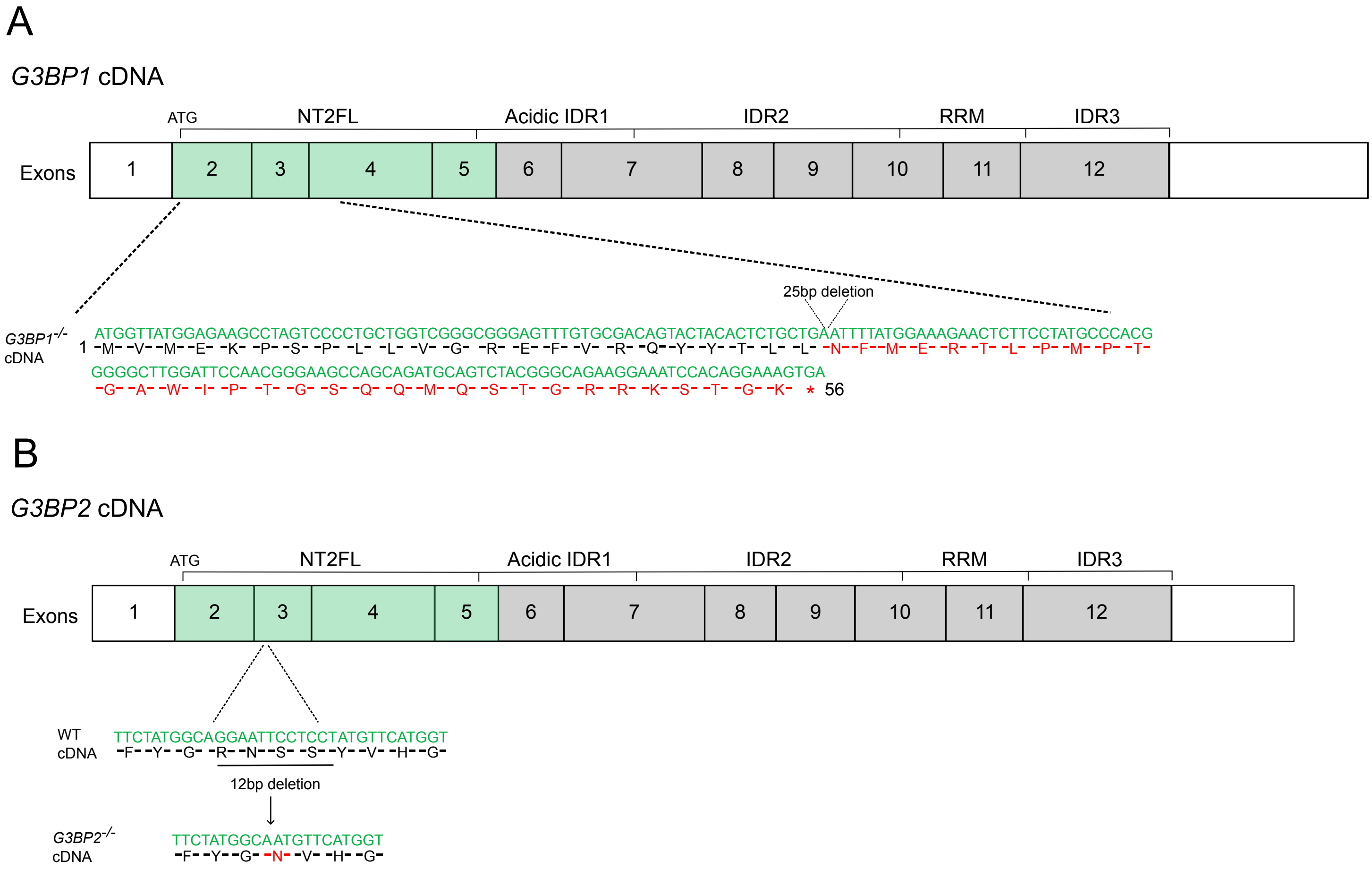
